# The output of interneurons in the primary visual cortex is best reflected by pre-synaptic activity, not somatic activity

**DOI:** 10.1101/2021.11.17.468122

**Authors:** Rozan Vroman, Lawrie McKay

**Affiliations:** Centre for Microsystems & Photonics, Department of Electronic and Electrical Engineering, University of Strathclyde, Glasgow, United Kingdom; Strategic Hub for Psychology, Social Work & Health Behaviours, School of Education & Social Science, University of the West of Scotland, Paisley, United Kingdom

**Keywords:** interneuron, pre-synapse, somatic activity, visual cortex, in vivo calcium imaging, vasoactive intestinal polypeptide, somatostatin, locomotion

## Abstract

Recent advances in 2-photon calcium-imaging in awake mice have made it possible to study the effect of different behavioural states on cortical circuitry. Many studies assume that somatic activity can be used as a measure for neuronal output. We set out to test the validity of this assumption by comparing somatic activity with the pre-synaptic activity of VIP (Vasoactive intestinal peptide)- and SST (Somatostatin)-positive interneurons in layer 2/3 of the primary visual cortex (V1). We used mice expressing genetically encoded calcium indicators in VIP/SST-interneurons across the whole cell (VIP/SST:GCaMP6f) or confined to pre-synapses (VIP/SST:SyGCaMP5). Mice were exposed to a full-field visual stimulation protocol consisting of 60-second-long presentations of moving Gabor gratings (0.04 cpd, 2 Hz) alternated by 30 seconds of grey screen. During imaging, mice were placed on an air-suspended Styrofoam ball, allowing them to run voluntarily. We compared neural activity during three 4-second time-windows: Before visual stimulation (−4 to 0 sec), during the initial onset (1 to 5 sec) and at the end of the stimulation (56 to 60 sec.). These were further compared while the mice were stationary and while they were voluntarily locomoting. Unlike VIP-somas, VIP-pre-synapses showed strong suppressive responses to the visual stimulus. Furthermore, VIP-somas were positively correlated with locomotion, whereas in VIP-synapses we observed a split between positive and negative correlations. In addition, a similar but weaker distinction was found between SST-somas and pre-synapses. The excitatory effect of locomotion in VIP-somas increased over the course of the visual stimulus but this property was only shared with the positively correlated VIP-pre-synapses. The remaining negatively correlated pre-synapses showed no relation to the overall activity of the Soma. Our results suggest that when making statements about the involvement of interneurons in V1 layer 2/3 circuitry it is crucial to measure from synaptic terminals as well as from somas.

## Introduction

Mapping the circuitry of micronetworks in sensory cortical areas is crucial for understanding the processing of sensory input and the integration of modulatory inputs. Neocortical neurons can be divided into excitatory pyramidal neurons and GABA-ergic inhibitory neurons, or interneurons. Despite pyramidal neurons accounting for about 80%-85% of cortical neuronal population (DeFelipe and Fariñas, 1992; Markram et al., 2004), classifying them into sub-types has proved hard (Kanari et al., 2019). Although interneurons compose a much smaller portion of the cortical neurons, they are crucial components in micro-networks and show high levels of diversity (Markram et al., 2004; Kepecs and Fishell, 2014). Interneuron classification based on gene expression has resulted in three main categories that cover most of the interneuron population and show limited overlap (Gonchar et al., 2008; Pfeffer et al., 2013). The three proteins upon which this is based are parvalbumin (PV), somatostatin (SST) and vasoactive intestinal peptide (VIP). Taniguchi et al. (2011) developed three interneuron cre-driver mouse lines based on the expression of these molecular markers and made it possible to map the connectivity patterns between them (Pfeffer et al., 2013). Furthermore, the interneuron cre-lines can be used to selectively express calcium indicators in the three cell types. Advances in 2-photon calcium imaging in awake and behaving mice (Dombeck et al., 2007; Goldey et al., 2014) improved our understanding of the interneuron circuitry involved in the processing of visual inputs and its modulation by changes in behavioural states such as locomotion (Fu et al., 2014; Dipoppa et al., 2018; Pakan et al., 2016; Millmann et al. 2020).

Most of these studies focus on somatic activity (e.g.: Fu et al. 2014; Karnani et al., 2016; Pakan et al., 2016; Jackson et al., 2016; Dipoppa et al., 2018; Millman et al., 2020) and in constructing connectivity maps based on their findings, make the assumption that somatic activity reflects a neuron’s output. However, calcium transients have been shown to vary along axons, dendritic spines and the soma (Koester and Sakmann, 2000; Francavilla et al., 2019), showing that changes in intracellular calcium concentrations are not homogeneous across a neuron.

Here, the assumption that somatic activity reflects the output of interneurons is tested for VIP and SST interneurons in the primary visual cortex V1 layer 2/3. These interneurons receive input from cholinergic neurons in the forebrain projecting to V1 and modulate visual processing (Picciotto et al., 2012; Adesnik et al., 2012; Alitto and Dan, 2013; Fu et al., 2014; Chen et al., 2015; Karnani et al., 2016; Pakan et al., 2016; Dipoppa et al., 2018; Millman et al., 2020). We compared somatic activity with pre-synaptic activity in response to prolonged full field visual stimulation, locomotion, and the interaction between them using 2-photon calcium imaging in awake behaving mice. Calcium indicators were either expressed throughout the neurons using GCaMP, or selectively in pre-synapses using SyGCaMP (Dreosti et al. 2009; Esposti et al. 2013; Baden et al. 2014; Schröder et al. 2019; Ryan et al., 2020). We confirmed that the somatic activity of VIP-neurons is positively correlated with locomotion (Fu et al., 2014; Pakan et al., 2016; Dipoppa et al., 2018; Reimer et al., 2014; de Vries et al., 2020), but that synapses show a split between positive and negative correlations. Furthermore, we found that whereas somatic activity does not show a significant response to visual stimulation, a pronounced suppressive response is observed at the pre-synaptic level. For SST-interneurons we found that both the somas and pre-synapses showed positive as well as negative correlations with locomotion (Pakan et al., 2016; Dipoppa et al., 2018). However, the negatively correlated synaptic boutons comprised a much smaller subset than the negatively correlated somas. Taken together, these results suggest that to draw conclusions on the output of cortical interneurons it is crucial to focus on pre-synaptic as well as somatic neural activity.

## Materials and Methods

### Animals

Cre-reporter mouse lines were used to confine the expression of genetically encoded calcium indicators to certain interneuron subtypes. The two cre-driver lines VIP-cre (*Vip*<tm1(cre)Zjh>/j) and SST-cre (*Sst*<tm2.1(cre)Zjh>/j) (Taniguchi et al., 2011) were cross-bred with the two reporter-lines GCaMP6f (129-S-Gt(ROSA)26Sor tm95.1 (CAG-GCaMP6f)) and SyGCaMP5 (Tg(ROSA26-LSL-SyGCaMP-WPRE)^L4^; Dreosti et al., 2009). The VIP-cre line was also used for virus injections with AAV9 phSyn1(S)-FLEX-SyGCaMP6f-WPRE acquired from VCF Charité (see section *Surgery and virus injection*). After allowing three days for recovery after surgery, mice were moved to a room equipped for housing under reversed light-dark conditions. Mice were provided with cage enrichments including scattered foraging food and a horizontal treadmill. To promote a good balance between locomotion and stationary periods during experiments, mice were well trained to be comfortable on the Styrofoam ball. Additional sessions of playtime with new toys and an upright exercise wheel were applied to further promote this balance.

Both males and females were used for experiments. All procedures were performed under personal and project licenses approved by the Animal Welfare and Ethical Review Body at the University of Sussex and released by the UK Home Office.

### Surgery and virus injection

Cranial window placement surgeries were performed on 8-12 week old mice. The surgical procedure was based on that from Goldey et al. (2014), but as this method was not fully aseptic, we had to adapt it to ensure it aligned with the guidelines of the Laboratory Animal Science Association (LASA). For a more detailed version see Boyd et al. (2021), but in brief:

Medication as per gram of mouse body weight: 4.8 μg dexamethasone, intramuscular injection 4-8 hours before surgery; 0.048 μg of the analgesic buprenorphine, intramuscular injection before or after surgery; 16 μl sterile saline, intraperitoneal injection after induction of general anaesthesia; 0.25 μg Metacam (meloxicam; nonsteroidal anti-inflammatory/analgesic), subcutaneous injection after induction of general anaesthesia; 0.4 μg Metacam mixed through wet mash food for 3 days after surgery.

General anaesthesia was induced by inhalation of a vapor comprising 4% isoflurane in medical oxygen. Based of animal’s body weight, 1-2% isoflurane was used to maintain anaesthesia. Body temperature was maintained at 37°C using a heating plate connected to a temperature controller (World Precision Instruments, USA). A rectal probe was used to monitor the animal’s body temperature. The mouse was placed in stereotax (Stoelting, US) and the eyes were kept moist by applying ophthalmic ointment (VitA-POS, Scope Ophthalmics, UK). Hair was removed around the surgical area, cleaned with saline and 70% alcohol, and an antiseptic was applied (betadine, Purdue Products, USA). From this point onward procedures were carried out using aseptic technique. Surfaces were cleaned with 70% alcohol and hands were washed with Hibiscrub (Centaur Services, UK) before putting on a sterile gown and sterile gloves. Surgical instruments (cleaned and autoclaved beforehand) were placed on sterile drapes. Skin was removed around an area above the left monocular V1 using spring scissors and the sides were sealed using Vetbond (3M, St Paul, US). Bleeding was minimised using spear sponges (Fine Science Tools, UK). The exposed skull was roughened using blunt forceps and overlapping scores in two perpendicular directions were made using a surgical scalpel blade. A surgical caliper was used to mark the centre of the craniotomy (3 mm lateral and 0.5 mm anterior of Lambda). A circle of 3 mm in diameter was drawn on the skull. To attach the custom-built titanium head post (see Goldey et al., 2014) dental cement (Unifast Trad Self Cure Liquid and powder, Kent Express Limited, UK) was used, mixed with a small amount of black tempera (1:15 ratio) for light shielding. The skull in the centre was left exposed. After the cement hardened, the mouse was placed in a free-moving anaesthesia mask and the head post was held in place using a lock-arm with an attached clamp. A fenestrated sterile drape was used to cover the mouse while leaving the surgical site exposed. A microdrill (Saeshin Precision Ind. Co., Korea) was used to drill a craniotomy over the marked circle and the surface around the circle was flattened to help the window attach later in the procedure. After soaking the bone with saline, a microprobe and/or forceps were used to lift off the bone flap and any resulting bleeding was controlled with gelfoam (Avitene Ultrafoam, Bard Davol Inc., USA). The dura was removed using superfine forceps (Dumont #5SF Forceps, Fine Science Tools). The glass window was constructed by gluing together three circular coverslips (a 5-mm coverslip with two 3-mm coverslips glued underneath; No. 1 thickness; Menzel-Glaser, Braunschweig, Germany) with UV-curing optical cohesive (NOI 61; Norland Products, Cranbury, NJ, USA). The window was cleaned with 70% ethanol and rinsed in saline before placing it on the craniotomy. The window was gently pushed down using an applicator attached to the stereotaxic manipulator arm and the top coverslip was attached to the skull using Vetbond. The window was further sealed using dental cement. Finally, two rubber rings were glued on top of the head post to create an imaging well as described by Goldey et al. (2014). By gradually lowering the percentage of isoflurane to 0%, the mouse was allowed to wake up. The mouse was kept warm and monitored until it started grooming and then placed back in its cage (it was made sure that the cage did not contain any parts that could cause the head post to get stuck).

In 3 mice, virus injections were performed during the surgery after removal of the bone-flap and dura. A microsyringe pump (UMP3 UltraMicroPump, Nanofil syringe; World Precision Instruments, Sarasota, FL, USA) was mounted on a digitalised stereotaxic arm (Stoelting, US). A bevelled borosilicate micropipette (GC100T-10, Harvard Apparatus) mounted on a Hamilton syringe was used for the injection of 200 nl AAV9 phSyn1(S)-FLEX-SyGCaMP6f-WPRE (titer: 2.38e12 GC/ml) in V1 at a depth of 250 μm.

### 2-Photon calcium imaging

Calcium imaging was performed on a Scientifica multi-photon scanning microscope using a Chameleon Ultra Diode-Pumped Laser and a wavelength of 940 nm. Images of 100 × 128 pixels were acquired at a rate of 10 Hz using the software package ScanImage r3.8.1. Mice were head-fixed under the objective and placed on an air-suspended Styrofoam ball. Locomotion was read out using an optical computer mouse (Dombeck et al., 2007) connected to an Arduino USB host shield/Arduino Uno motherboard, which was in turn connected to the acquisition computer (Dell, Windows 7). PsychoPy, a stimulation toolbox library for Python, was used to generate the visual stimulation on a CyberPowerPC running Ubuntu. Two 535 × 300 mm screens (BenQ, XL2411Z 24 inch) were used for stimulus presentation (framerate: 60Hz). Each screen spanned 184° of the horizontal visual field and 105° of the vertical visual field. Signals for the on and offset of the visual stimulus were sent to the acquisition computer using a U12 Labjack.

### Visual stimulation protocol

Drifting Gabor gratings at an angle of 315° and a temporal and spatial frequency of 2 Hz and 0.04 cycles per degree respectively were used as visual stimulation. They were presented for periods of 60 seconds and followed by a period of grey screen with equal luminance. As pilot data suggested that it may take just over 20 seconds for the baseline to recover after visual stimulation (Supplementary Figure 3A & B), the duration of grey screen was set at 30 seconds. The full protocol for one experiment consisted of 5 seconds of grey screen followed by 5-10 repetitions of the stimulation sequence (60 sec. of Gabor gratings + 30 seconds of grey screen); figure 3B shows a section of a timeseries using the protocol. At the start of each session, before starting the experiments, the mice were presented with a series of visual stimulations. The reason for this was the finding that when the mice were not exposed to the visual stimulation for at least 24 hours, the response to the first stimulus presentation was consistently excitatory instead of suppressive (Supplementary Figure 2). Well-trained mice were exposed to experimental sessions of up to two hours.

### Image analysis

Most of the analysis was performed using Igor Pro 6.36 and the Sarfia plug-in (Dorostkar et al, 2010). Image stacks were first registered using the inbuilt registration function from Igor pro to correct for x-y drift. Regions of interest (ROIs) were generated using the thresholding tool from Sarfia and was based on the standard deviation map of the neuronal activity. The results were manually adjusted if necessary. ROIs that were touching the borders of the field of view were sometimes affected by the registration procedure, leading to artefacts. These ROIs were removed before further analysis. Further adjustments were performed to make sure every ROI had a diameter of 2-3 pixels, equal to 0.7-2.1 μm for the VIP-synapses and 0.6-1.7 μm for the SST-synapses. The baseline fluorescence F_0_ was determined by averaging the neural activity without locomotion or visual stimulation early during the recording. There was one exception in that for the VIP-soma recordings under the Locomotion condition the baseline needed to be taken during locomotion. ΔF/F_0_ was calculated using the formula ΔF/F_0_ = (F-F_0_)/F_0_ where F is the absolute fluorescence.

A set of criteria was used to determine if a dataset was included in further analysis: (1) As bouts of locomotion and stationary periods needed to be present during both grey screen and visual stimulation, any dataset without this was excluded. (2) If the registration to correct for x-y drift failed at any point during the experiment, the dataset was excluded. (3) If some or all synapses showed drops in signal the dataset was excluded. (4) If a slow drift in baseline activity was detected, the dataset was excluded.

### PCA correction method

Principal component analysis (PCA) was applied to the data collected from VIP-and SST-synapses using Python 7.2 (sklearn.decomposition.PCA). Igor pro 6.36 was used to calculate the delay with locomotion of all components. Because the response to locomotion is a multisynaptic event, most synpases showed a delay of around 300 ms (supplementary figure 1A). Components with a delay of 0 ± 100 ms were regarded as physiologically infeasible and therefore removed, and the data was then reconstructed using Python 2.7. See Supplementary Figure 1 for a detailed description of the method. All further analysis was conducted on the corrected data.

### Correlation and bootstrapping method

Correlations coefficient values of the calcium imaging traces with the locomotion time series were calculated using Spearman’s rank correlations due to the non-normality of the imaging data. To determine the probability that findings were of real effects (p-value) an adaptation of the shuffling method described by Dipoppa et al. (2016) was used. In this bootstrapping method, a distribution of correlation coefficients is generated using a 1000 random circular shifts of the locomotion speed trace relative to the fluorescence trace, with the p-value derived as the probability the data is from one of these random distributions. In our procedure, the locomotion trace (10 Hz) was first smoothed by 5 points and a list of p-values for every ROI within a field of view was generated using the shuffling method. This list was then corrected for multiple comparison using the Benjamini and Hochberg False Discovery Rate method (Benjamini and Hochberg, 1995). For grouping the pre-synapses into the Positive and Negative group, ROIs with significant (p<0.05 after FDR correction) positive correlation coefficient values were placed in the Positive group and ROIs with significant negative values were placed in the Negative group.

### Statistics

Each experiment consisted of a recording from a field of view of 240 μm × 190 μm in the case of somatic recordings and 66 μm × 66 μm in the case of recordings from pre-synaptic boutons, during the presentation of the visual stimulation protocol (a total duration of 8-15 min. per experiment). For each experiment, the data obtained from the ROIs were averaged. In the case of the synapses however, these were split into the Positive and Negative groups before averaging. As a result, the sample size was equal to the number of experiments. This was done because the neural responses within the same experiment are correlated and thus combining the ROIs prevents incorrect inflation of the degrees of freedom (Galbraith et al., 2010).

The data from each experiment was subsequently split into two Locomotion conditions: Locomotion and Stationary. Data recorded during each visual simulation window were extracted along with 4 seconds of grey screen prior to the stimulus onset to use as a baseline. This resulted in between five and ten 64s timeseries for each experiment, from which the mean timeseries was calculated. There were 3 parts of the timeseries of interest – baseline (−4-0s), activity during early onset (1-5s) and activity at the end of stimulus presentation (56-60s), which we refer to from now on as T1, T2 and T3 respectively; these are indicated in the figures by respectively red, green and blue bars (Figures 1D,E, 3C,D, 5C,D and 7). Data from these windows were extracted from the mean timeseries, then the mean was calculated for each. This data extraction was calculated for all conditions.

**Figure 1.**
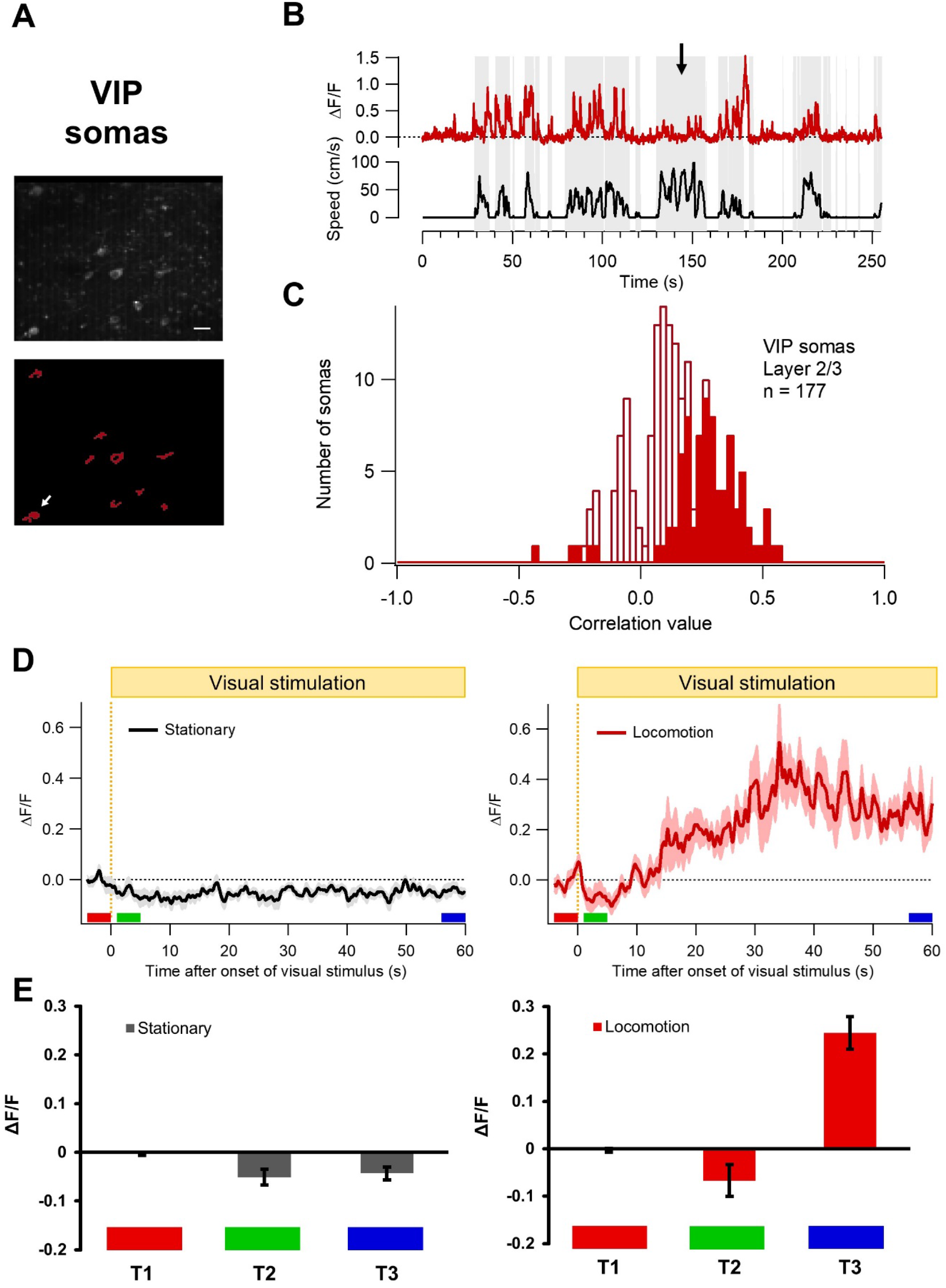
The effect of locomotion and prolonged visual stimulation on the somatic activity of VIP-positive interneurons in V1 layer 2/3. **(A)** An example of a field of view (top) and the ROIs corresponding to the somas (bottom). Scale bar represents 20 μm. **(B)** An example of a fluorescence timeseries extracted from the ROI indicated in **A** by the white arrow. The red trace represents the somatic activity with time in seconds on the x-axis and ΔF/F_0_ on the y-axis. The black trace represents the locomotion speed in cm/s. The arrow indicates an example where a bout of fast running initiates only minor increases in activity, showing that whereas locomotion and somatic activity are strongly correlated, some variability can be found. **(C)** Histogram of somas and their correlation value with locomotion. The filled bars represent correlations with p-values < 0.05 after FDR correction. The total number of somas was 177, recorded from 3 mice. **(D)** The average somatic activity before and during prolonged visual stimulation. The x-axis represents the time relative to the onset of the visual stimulus and the y-axis represents the fluorescence in ΔF/F_0_. The black trace represents the somatic activity while the mouse is stationary and the red trace while the mouse is in locomotion. The light area around the traces represents the standard error of the means (SEM). **(E)** The average activity was measured at three timepoints (indicated in **D**): before onset of visual stimulation (T1, red, −4-0 sec.), during early visual stimulation (T2, green, 1-5 sec.) and after almost a minute of visual stimulation (T3, blue, 56-60 sec.). Left: stationary, right: locomotion. Error bars represent SEM.

To determine whether there was a significant interaction between visual stimulation and locomotion in VIP-somas (3 mice), we utilised a 2×3 mixed design with a within-subject factor of Timepoint with 3 levels (T1, T2 and T3) and a between-subjects factor of Locomotion condition with 2 levels (Locomotion and Stationary).

To determine whether there were significant interactions between visual stimulation and locomotion in VIP-synapses (crossbred mice (5) and virus injected mice (3)) and SST-synapses (3 mice), we utilised a 2×3×2 mixed design with 2 within-subjects factors: Locomotion condition with 2 levels (Locomotion and Stationary) and Timepoint with 3 levels (T1, T2 and T3). The between-subjects factor was Correlation Sign with 2 levels (Positive group and Negative group). The dependent variable is the mean neural activity as measured by fluorescence in ΔF/F_0_.

All data was tested for sphericity using the Greenhouse-Geisser test and the assumption was not found to have been violated.

Mixed ANOVA analyses and t-tests were performed using IBM SPSS Statistics, and follow up Wilcoxon matched-pairs Signed-Rank Tests were performed using R studio.

### Results

In this study we set out to compare the somatic activity of VIP-interneurons and SST-interneurons with pre-synaptic activity in mouse V1, layer 2/3. We used genetically encoded calcium indicators and 2-photon fluorescence imaging. Mice from the VIP-cre and SST-cre driver lines (Taniguchi et al., 2011) were crossed with either the GCaMP6f or SyGCaMP5 reporter lines in order to perform 2-photon imaging of, respectively, interneuron somas and pre-synaptic boutons in awake, behaving mice. Removable cranial windows were placed over V1 in the left hemisphere of the mice (Goldey et al., 2014). Mice were head-restrained under the objective lens of a 2-photon microscope and placed on an air-suspended Styrofoam ball (Dombeck et al., 2007). An optical computer mouse placed close to the ball was used to read out the locomotion (Dombeck et al., 2007).

### Neural activity of VIP-somas in relation to locomotion and visual stimulation

#### VIP-somas are positively correlated with locomotion

Our first point of focus was the effect of locomotion on somatic activity of VIP-interneurons in the absence of visual stimulation (grey screen). Figure 1A shows an example of a field of view (standard deviation map) and the corresponding regions of interest (ROIs) of VIP-somas from a VIP:GCaMP6f mouse. Figure 1B shows the fluorescence timeseries (red trace) extracted from the ROI indicated by the white arrow in the lower panel of Figure 1A. Fluorescence traces representing neural activity were expressed as relative changes in fluorescence by dividing the change in fluorescence by the baseline fluorescence early in the recording (ΔF/F_0_). The black trace depicts the locomotion in cm/s (Figure 1B). A clear correlation between the somatic activity and locomotion was observed with bouts of locomotion eliciting strong increases in neural activity, albeit with some exceptions that only elicited a moderate increase (Figure 1B, arrow). To quantify this the Spearman correlation coefficient values were calculated for all somas and plotted as a histogram (Figure 1C). The red filled bars (Figure 1C) represent the values that were significantly correlated with locomotion. The VIP-somas are predominantly positively correlated (140 out of 177, of which 78 were significant; p<0.05; N=3) with only a small group of negatively correlated somas (37 out of 177, of which 5 were significant; p<0.05; N=3). This is in line with findings from previous studies (Dipoppa et al., 2018; Pakan et al., 2016; Fu et al., 2014; Reimer et al., 2014).

#### The interaction of locomotion and visual stimulation

Next, we set out to investigate the interaction of locomotion and visual stimulation in the VIP-somas. The visual stimulus we used was a full field drifting Gabor grating at an angle of 315° (to prevent interaction with processing related to optic flow), a contrast of 1, a spatial frequency of 0.04 cycles per degree and a temporal frequency of 2 Hz. With the aim to study both initial responses to the visual stimulus and responses after prolonged exposure, each stimulus presentation lasted for 60 seconds. The full protocol started with 5 seconds of homogeneous illumination (grey screen) followed by five 60-second stimulations that were interleaved with 30 seconds of grey screen. The timeseries of the five repetitions were averaged as described in the Materials and Methods. We collected data from 3 mice under two conditions of Locomotion: the Stationary condition during which mice were sitting on the Styrofoam ball and the Locomotion condition during which mice were voluntarily locomoting. From each experiment, we extracted 3 Timepoints as described in the Materials and Methods section: T1 right before visual stimulation, T2 after the onset of visual stimulation with a 1-second offset and T3 right before the end of the visual stimulation.

Pilot data from VIP-somas using 5-second episodes of visual stimulation suggested a small suppression in response to full field visual stimulation (Supplementary Figure 3A). This has indeed been reported in literature (de Vries et al. 2020; Millman et al., 2020), although Pakan et al. (2016) found a small but non-significant suppression in neural activity. Pakan et al. (2016) show excitatory responses to locomotion both without visual stimulation (in their case darkness) and during prolonged visual stimulation. Considering these findings, we hypothesised that during the Stationary condition we would find a suppression of neural activity during Timepoints 2 and 3, but no response to visual stimulation in the Locomotion condition.

Figure 1D shows the mean somatic responses for the Stationary condition (left) and the Locomotion condition (right). The red, green and blue blocks indicate the three timepoints. In Figure 1E the means for each timepoint are plotted as a bar graph with standard error of the means for each Locomotion condition (Stationary on the left, Locomotion on the right). A small suppression of neural activity was observed from Timepoint 1 to Timepoint 2 and 3 in response to the visual stimulus for the Stationary condition (T1: ΔF/F_0_ of −0.003 ± 0.003, T2: ΔF/F_0_ of −0.051 ± 0.016, T3: ΔF/F_0_ of −0.043 ± 0.013). A similar reduction in the initial response to the visual stimulus was also observed for the Locomotion condition (T1: ΔF/F_0_ of −0.004 ± 0.004, T2: ΔF/F_0_ of −0.067 ± 0.034). However, the average neural activity at timepoint 3 was higher than the baseline level (ΔF/F_0_ of 0.244 ± 0.034).

#### VIP-somas show an increasingly excitatory response to prolonged visual stimulation during locomotion

To test our initial hypotheses, we performed a 2×3 mixed ANOVA. There was a significant main effect of both Time F(2,22)=25.344, p=0.000002, n^2^_p_=0.697, Power=1 and Locomotion condition F(1,11) = 28.264, p=0.000246, n^2^_p_=0.720, Power = 0.998. The main effects were, however, confounded by a significant interaction between Time and Locomotion F(2,22)=28.169, p=8.569.10^−7^, n^2^_p_ =0.719, Power=1.

To further elucidate the interaction, simple main effects were explored using a one-way ANOVA for each Locomotion condition. The one-way ANOVA for the Stationary condition found a significant effect of Time F(2,12)=7.737, p=0.007, n^2^_p_=0.563, Power=0.880. However, post-hoc tests could not determine where the significant differences lay. We can therefore not conclusively determine whether the visual stimulation resulted in a suppressive response and therefore have to reject the hypothesis of finding a suppression of neural activity in the Stationary condition at Timepoints 1 and 2. In the Locomotion condition there was a significant main effect of Time F(2,10)=24.473, p=0.000141, n^2^_p_=0.830, Power=1. Post-hoc tests revealed the main effect of Time was driven by a significantly higher level of fluorescence at timepoint 3 than either timepoint 1 (mean difference=0.243, p=0.005) or timepoint 2 (mean difference=0.307, p=0.012). This suggests that for the Locomotion condition there was no significant initial response to visual stimulation, but that after prolonged visual stimulation neural activity is significantly increased (Figure 1D), which is in line with the findings of Pakan et al. (2016). Based on these results we can accept the hypothesis that a suppressive response to visual stimulation is absent in the Locomotion condition.

### Neural activity of VIP-synapses in relation to locomotion and visual stimulation

#### Correction for drift-artefacts

With the aim to compare somatic with pre-synaptic activity, we repeated the experiments in 5 VIP:SyGCaMP5 mice that had a calcium indicator selectively expressed in pre-synaptic boutons (Dreosti et al., 2009). As synaptic boutons are considerably smaller than somas (<1 μm), measuring changes in fluorescence in boutons are much more prone to drift-artefacts (Ryan et al., 2020). Whereas drift in the x- and y-direction, mainly caused by locomotion, can be corrected for with registration, correction for the z-direction is trickier. Ryan et al. (2020) developed a method to correct for z-motion artefacts by measuring displacements using the additional imaging of blood vessels as an anatomical marker. As the data presented here was collected before this method was developed, a different approach was used. Firstly, data was included based on a set of criteria (see methods) to make sure the analysis was performed on a high-quality dataset. Secondly, we developed and applied a method based on principal component analysis, a statistical tool commonly used for timeseries analyses in neuroscience studies, (e.g fMRI and EEG) to deal with motion artefacts. See the Methods section and Supplementary Figure 1 for a detailed description. All data collected from synapses was corrected in this way before further analysis.

#### Experimental design

To allow for comparisons of responses from the same synapse under all conditions (visual stimulation vs grey screen during locomotion and stationary periods), we used a prolonged visual stimulation protocol (60 sec. visual stimulation/30 sec. grey screen; 5-10 repetitions; full field drifting Gabor grating, 315°, contrast=1, spatial frequency=0.04 cpd, temporal frequency=2Hz) and only timeseries that included bouts of voluntary locomotion and stationary periods during both visual stimulation and grey screen were included in the analysis. In this way, the Locomotion condition (Locomotion and Stationary) could be treated as a within subjects independent variable.

#### VIP-synapses show both positive and negative correlations with locomotion

First, we calculated the correlation of the synaptic activity of VIP-interneurons with locomotion during grey screen. Figure 2A shows an example of a field of view and the corresponding ROIs below. Interestingly, both positive and negative correlations were found, with 340 out of 1103 pre-synaptic boutons showing a significant negative correlation (p<0.05; N=5) and 224 showing a significant positive correlation with locomotion (p<0.05; N=5; Figure 2C). This is in line with the findings of Ryan et al. (2020) and reveals a clear discrepancy between somatic and synaptic activity in relation to locomotion. Figure 2B shows an example of a fluorescence timeseries for both a synapse that is positively correlated with locomotion (yellow trace) and a synapse that is negatively correlated (blue trace; the black trace represents the speed of locomotion; the two example synapses are indicated by a white arrow in Figure 2A).

**Figure 2.**
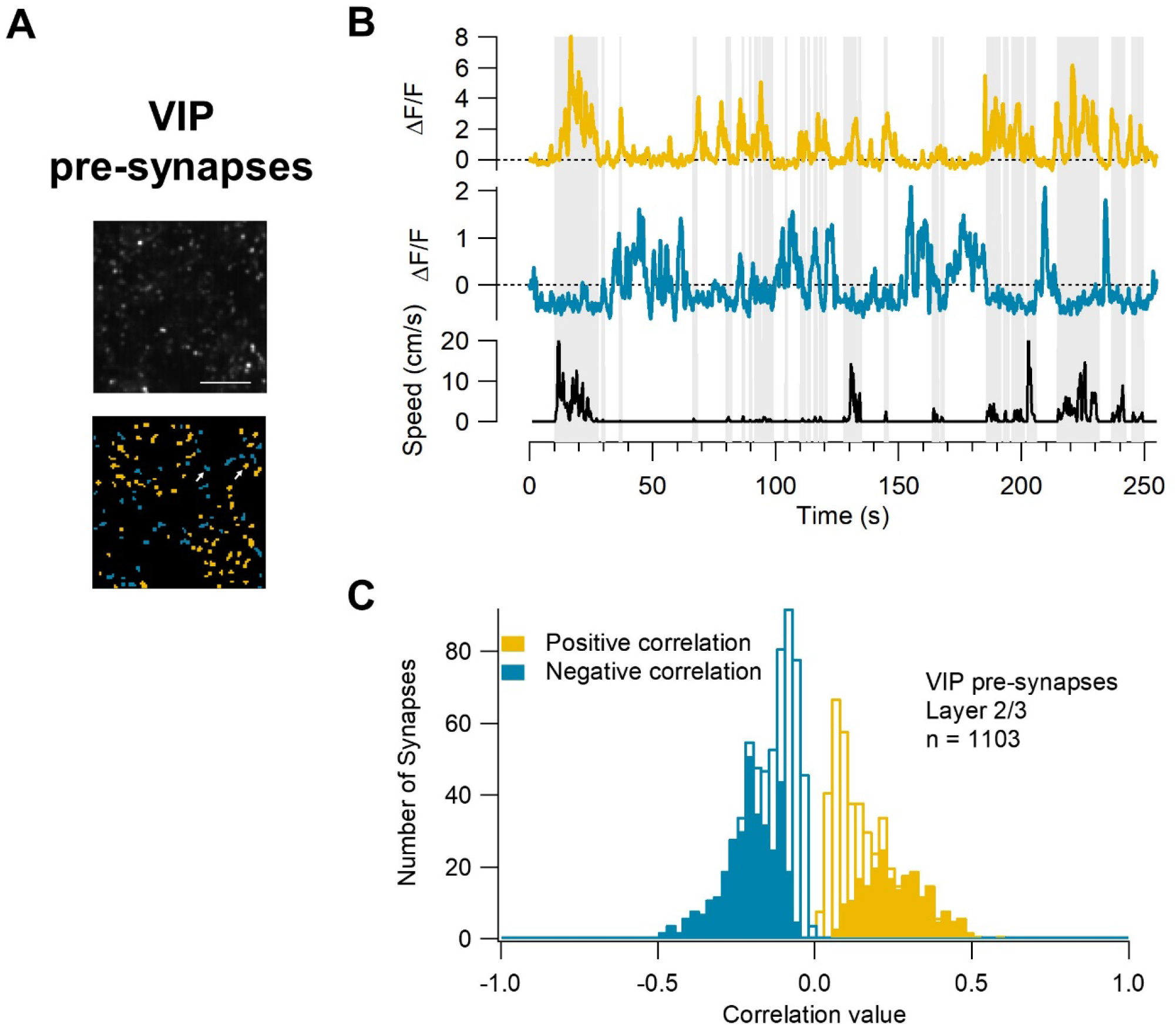
The effect of locomotion on the pre-synaptic activity of VIP-positive interneurons in V1 layer 2/3. **(A)** An example of a field of view (top) and the ROIs corresponding to the synapses (bottom). The ROIs showing a positive correlation with locomotion are in yellow and the ROIs showing negative correlations are in blue. Scale bar represents 20 μm. **(B)** An example of a fluorescence timeseries extracted from a ROI with a positive correlation with locomotion (yellow timeseries) and a ROI with a negative correlation (blue timeseries). The two ROIs are indicated in **A** by white arrows. The black trace represents the locomotion speed in cm/s. **(C)** Histogram of VIP-synapses and their correlation value with locomotion. The filled bars represent correlations with p-values < 0.05 after FDR correction. The total number of synapses was 1103, recorded from 5 mice.

#### The interaction of locomotion and visual stimulation

Next, we set out to investigate the interaction of locomotion and visual stimulation in the VIP-synapses (Figure 3). Preliminary data suggested that there was a suppressive response to full field visual stimuli with the spatial and temporal frequencies of 0.04 cpd and 2 Hz (Supplementary Figure 3B). Furthermore, preliminary data also suggested that this was the case for the Locomotion condition (Supplementary Figure 3C). We therefore hypothesised that, unlike the VIP-somas, VIP-synapses would have a significant initial suppression in response to visual stimulation both for the Stationary and Locomotion conditions. We further hypothesised that for the Stationary condition this suppression would be maintained, whereas for the Locomotion condition synaptic activity would become increasingly excitatory as was observed in the soma.

**Figure 3.**
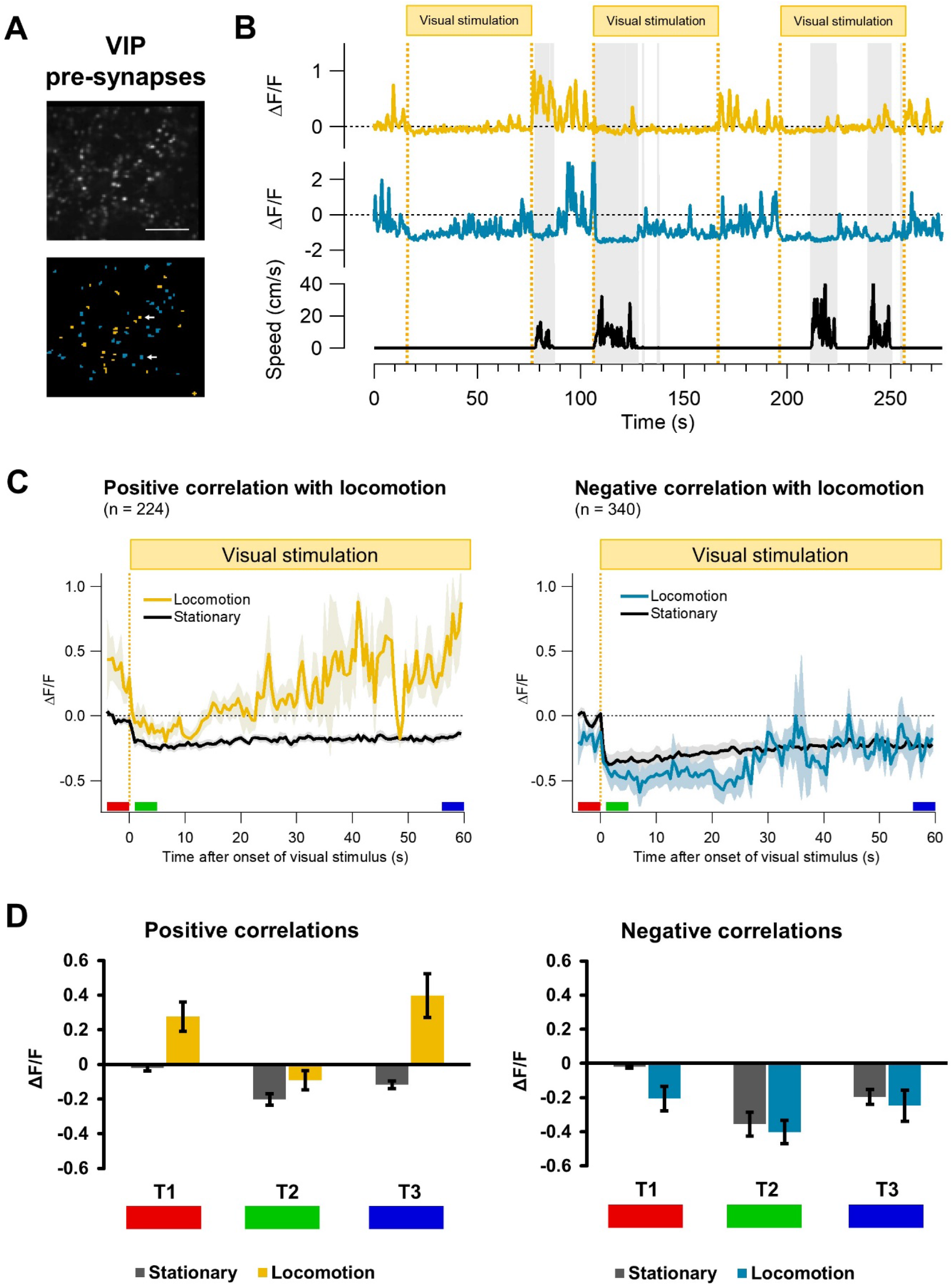
The effect of prolonged visual stimulation, with and without locomotion, on the pre-synaptic activity of VIP-positive interneurons in V1 layer 2/3. **(A)** An example of a field of view (top) and the ROIs corresponding to the synapses (bottom). The ROIs showing a positive correlation with locomotion are in yellow and the ROIs showing negative correlations are in blue. Scale bar represents 20 μm. **(B)** An example of a fluorescence timeseries extracted from a ROI with a positive correlation with locomotion (yellow timeseries) and a ROI with a negative correlation (blue timeseries). The visual stimulation periods are indicated at the top, with the yellow dotted lines indicating the start and end of each 60 second presentation. The two ROIs are indicated in **A** by white arrows. The black trace represents the locomotion speed in cm/s. **(C)** The average synaptic activity before and during prolonged visual stimulation. Synapses were separated into a group containing synapses that showed significant positive correlations with locomotion (Positive group) and a group representing significant negative correlations (Negative group). The x-axis represents the time relative to the onset of the visual stimulus and the y-axis represents the fluorescence in ΔF/F_0_. The black trace represents the synaptic activity while the mouse is stationary and the yellow (left; Positive group) and blue traces (right; Negative group) while the mouse is in locomotion. The light area around the traces represents SEM. **(D)** The average activity was measured at three timepoints (indicated in **C**): before onset of visual stimulation (T1, red), during early visual stimulation (T2, green) and after almost a minute of visual stimulation (T3, blue). The dark grey bars represent neural activity during stationary periods and the yellow (left) and blue (right) bars activity during periods of locomotion. Error bars represent SEM.

Figure 3A shows an example field of view with the corresponding ROIs. The timeseries of the two synapses indicated by white arrows are shown in Figure 3B. The data was split into two Correlation groups: ROIs showing significant positive correlations with locomotion (Positive group) and ROIs showing significant negative correlations (Negative group). Figure 3C shows the average responses of each group to the visual stimulation while the mouse is stationary (black trace) and while the mouse is locomoting (yellow trace and blue trace for the Positive and Negative group respectively). For further analysis, the average of three 4-second time windows were calculated as before: T1 right before visual stimulation, T2 after the onset of visual stimulation with a 1-second offset and T3 right before the end of the visual stimulation (Figure 3C,D). In Figure 3D these values are plotted in a bar graph for the Positive group (left) and the Negative group (right). For both the Positive and Negative group the visual stimulation resulted in a suppressive response when the mouse was stationary (Positive group: T1: −0.020 ± 0.019 ΔF/F_0_, T2: −0.203 ± 0.032 ΔF/F_0_, T3: −0.117 ± 0.022 ΔF/F_0_; Negative group: T1: −0.019 ± 0.009 ΔF/F_0_, T2: −0.355 ± 0.070 ΔF/F_0_, T3: −0.196 ± 0.044 ΔF/F_0_). Note that these responses are much more pronounced in the synapses than in the soma. When the mouse was locomoting, the Positive group showed a suppression in neural activity early in the visual stimulation followed by a gradual increase over the course of the minute-long presentation (T1: 0.276 ± 0.085 ΔF/F_0_, T2: − 0.092 ± 0.055 ΔF/F_0_, T3: 0.398 ± 0.126 ΔF/F_0_). The neural activity during the visual stimulation in the Negative group showed a similar pattern, but average fluorescence levels were generally lower. Furthermore, the average levels of fluorescence remained below baseline levels throughout the visual stimulation and did not differ much from the Stationary condition (T1: −0.206 ± 0.072 ΔF/F_0_, T2: −0.402 ± 0.068 ΔF/F_0_, T3: −0.248 ± 0.091 ΔF/F_0_).

#### VIP-synapses show a suppressive initial response to visual stimulation in all conditions

To test for significant differences between the groups and conditions, we performed a 2×3×2 mixed ANOVA. There was a significant main effect of Locomotion F(1,17)=6.16, p=0.024, n^2^- _p_=0.266, Power=0.648, Time F(2, 34)=17.63, p=0.000006, n^2^_p_=0.509, Power=1.000 and Correlation group F(1,17)=9.42, p=0.007, n^2^_p_=0.357, Power=0.824. Post-hoc tests revealed that the main effect of Time was driven by a significantly lower level of fluorescence at timepoint 2 than either timepoint 1 (mean difference=-0.283, p=0.000238) or timepoint 3 (mean difference=-0.1.99, p=0.001). Timepoint 1 and 3 did not differ significantly from each other (mean difference=0.084, p=0.279).

There was no interaction between Time and Correlation group F(2,34) = 17.634, p=0.872, n^2^- _p_=0.008, Power=0.069 or between Locomotion and Time F(2,17)=3.005, p=0.063, n^2^_p_=0.150, Power=0.545, and no three-way interaction F(2,34)=2.253, p=0.121, n^2^_p_=0.117, Power=0.427. However, the main effects of Locomotion and Correlation group were confounded by a significant interaction between Locomotion and Correlation group F(1,17)=21.48, p=0.000237, n^2^_p_=0.558, Power=0.992. To determine whether each Correlation group differed significantly from each other, independent samples T-tests were conducted separately for each Locomotion condition. Equality of variance was evaluated using Levene’s Test, which found that the assumption was violated only for the Stationary condition (F=10.299, p=0,003), requiring the degrees of freedom to be adjusted. The Correlation groups did not differ in the Stationary condition t(20.508)=1.842, p=0.080, but mean fluorescence levels were found to be significantly higher in the Positive group than the Negative group t(25)=5.644, p=0.000007.

To further test whether the levels of neural activity differed from baseline levels for the Locomotion conditions and Correlation groups in response to visual stimulation at timepoints 2 and 3, we performed a one-sample t-tests against 0. As we performed 12 tests in total, the Benjamini and Hochberg False Discovery Rate method (Benjamini and Hochberg, 1995) was used to correct for multiple comparisons. Both the original p-values and the adjusted values (q-values) are reported. For the Stationary condition we found that the fluorescence level at timepoint 2 was significantly below baseline levels in both the Positive group (mean difference=-0.203, p=0.000017 q=0.000207) and the Negative group (mean difference=-0.355, p=0.000159, q=000476). This was similar at timepoint 3 for both the Positive group (mean difference=-0.117, p=0.000092, q=0.000368) and the Negative group (mean difference=-0.196, p=0.000575, q=0.00138), confirming the hypothesis that in the Stationary condition there is a significant suppressive response to the visual stimulation. However, it must be noted that this suppression decreased over the course of the visual stimulation, confirmed by the significant main effect of time reported above.

#### VIP-synapses in the Positive group show an increasingly excitatory effect of locomotion in response to prolonged visual stimulation

In the Locomotion condition, both the Positive and the Negative group showed a decrease in neural activity at timepoint 2, but only in the Negative group was the level of fluorescence significantly below 0 (Positive group: mean difference=-0.092, p=0.120 q=0.131; Negative group: mean difference=0.402, p=0.000053, q=0.000317). In the Negative group the average fluorescence level at timepoint 3 in the Locomotion condition was significantly below 0 (mean difference=0.248, p=0.018, q=0.024) and did not exceed the average level in the Stationary condition. As a result, the hypothesis that locomotion has an increasingly excitatory effect after prolonged visuals stimulation has to be rejected for the Negative group. The average fluorescence level for the Positive group in the Locomotion condition at timepoint 3 was significantly above 0 (mean difference=0.398, p=0.0093, q=0.019). As such, the hypothesis that after prolonged visual stimulation locomotion has an excitatory effect can be confirmed for the Positive group only, even though this effect was less pronounced than in the somas.

### SST-somas showed both positive and negative correlations with locomotion

Next, we measured neural activity in somas of SST-positive neurons in 3 SST:GCaMP6f mice and calculated the Spearman correlation with locomotion (Figure 4C). SST somas showed both positive and negative correlations with locomotion; 29 out of 114 somas showed a significant negative correlation and 37 a significant positive correlation (N=3; Figure 4C). Figure 4A shows an example of a field of view and the corresponding ROIs. The white arrows indicate an example of a soma showing a positive and a negative correlation with locomotion. In Figure 4B a section of the corresponding time-series is shown (yellow=positive correlation; blue=negative correlation; black=locomotion).

**Figure 4.**
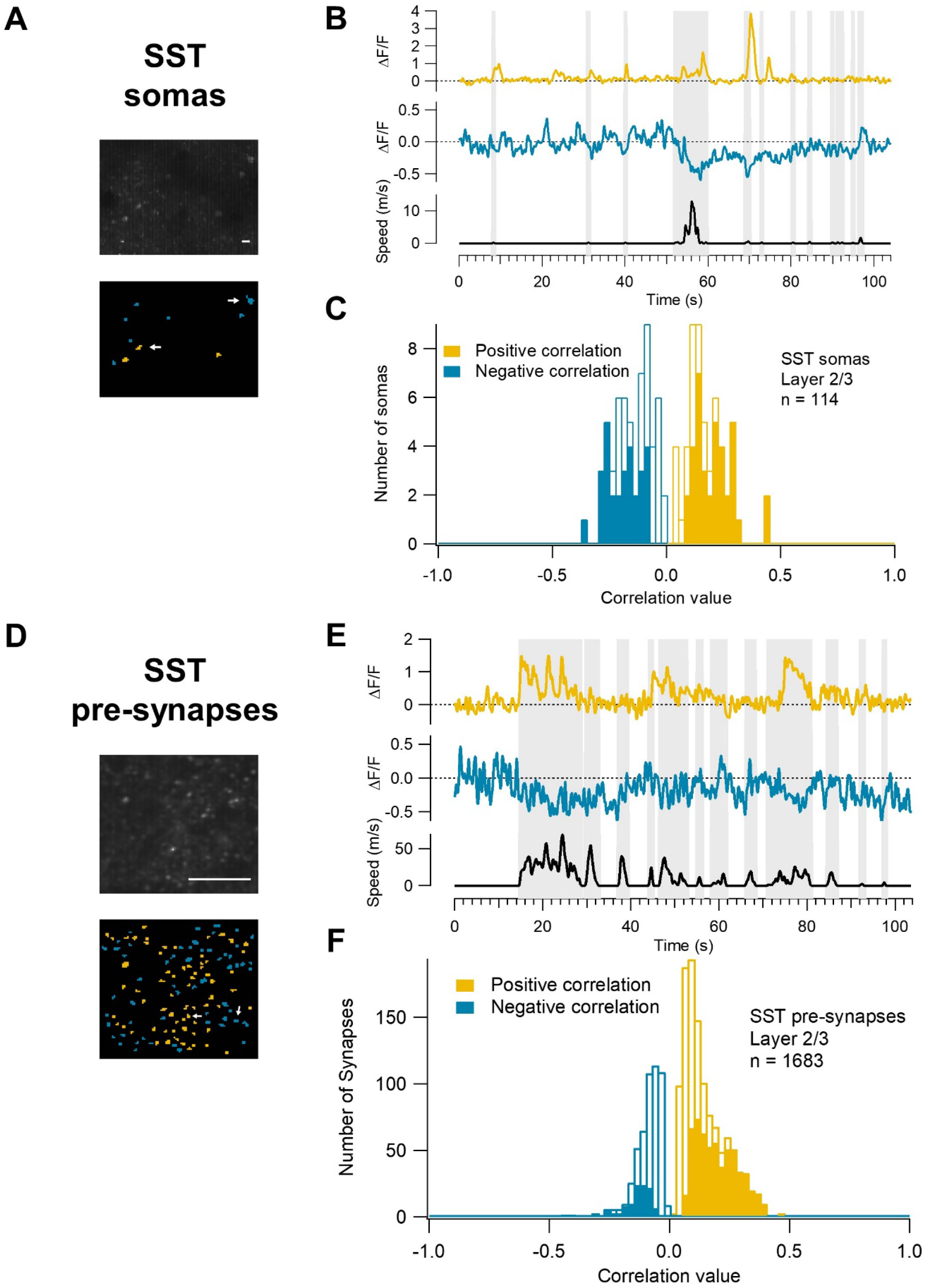
The effect of locomotion on the somatic and pre-synaptic activity of SST-positive interneurons in V1 layer 2/3. **(A)** An example of a field of view (top) and the ROIs corresponding to the somas (bottom). The ROIs showing a positive correlation with locomotion are in yellow and the ROIs showing negative correlations are in blue. Scale bar represents 20 μm. **(B)** An example of a fluorescence timeseries extracted from a ROI with a positive correlation with locomotion (yellow timeseries) and a ROI with a negative correlation (blue timeseries). The two ROIs are indicated in **A** by white arrows. The black trace represents the locomotion speed in cm/s. **(C)** Histogram of SST-somas and their correlation value with locomotion. The filled bars represent correlations with p-values < 0.05 after FDR correction. The total number of somas was 114, recorded from 3 mice. **(D)** An example of a field of view (top) and the ROIs corresponding to the synapses (bottom). The ROIs showing a positive correlation with locomotion are in yellow and the ROIs showing negative correlations are in blue. Scale bar represents 20 μm. **(E)** An example of a fluorescence timeseries extracted from a ROI with a positive correlation with locomotion (yellow timeseries) and a ROI with a negative correlation (blue timeseries). The two ROIs are indicated in **D** by white arrows. The black trace represents the locomotion speed in cm/s. **(F)** Histogram of SST-synapses and their correlation value with locomotion. The filled bars represent correlations with p-values < 0.05 after FDR correction. The total number of synapses was 1683, recorded from 3 mice.

### Neural activity of SST-synapses in relation to locomotion and visual stimulation

To compare the somatic activity of SST-positive neurons with the pre-synaptic activity we recorded from 3 SST:SyGCaMP5 mice. The same 2×3×2 mixed design was used as for the VIP-synapses and the same visual stimulation protocol was used (60 sec. visual stimulation/30 sec. grey screen; 5-10 repetitions; full field drifting Gabor grating, 315°, contrast=1, spatial frequency=0.04 cpd, temporal frequency=2Hz).

#### A small minority of SST-synapses are negatively correlated with locomotion

First, we calculated the Spearman correlations of the synaptic activity with locomotion (Figure 4F). Again, both positive and negative correlations were found. Interestingly, however, the group of negatively correlated SST-synapses was much smaller than the Positive group. 118 out of 1683 showed a significant negative correlation with locomotion while 593 were significantly positively correlated (N=3; Figure 4F). Figure 4D shows an example of a field of view with the corresponding ROIs. A section of the time-series corresponding to the two synapses indicated by white arrows are shown in Figure 4E (yellow and blue traces; black trace represents locomotion).

#### The interaction of locomotion and visual stimulation

Next, we addressed SST-synaptic responses to visual stimulation and the effect of locomotion. Pakan et al. (2016) report that SST-somas show excitatory responses to visual stimulation. Furthermore, they show that whereas in the dark they find both positive (24 ± 6%) and negative (11 ± 5%) correlations with locomotion, during visual stimulation they found many more positive (63 ± 7%) than negative correlations (4 ± 3%). We therefore hypothesised that both Correlation groups would show a significant excitatory response to visual stimulation, both at timepoint 2 and 3. We further hypothesised that in the Locomotion condition the level of fluorescence at these timepoints would be significantly higher than in the Stationary condition for both the Positive and Negative group.

Figure 5A shows an example field of view with the corresponding ROIs. The timeseries of the two synapses indicated by white arrows are shown in Figure 5B. As with the VIP-synapses, the data was split into two Correlation groups: the Positive and the Negative group. Figure 5C shows the average responses of each group to the visual stimulation while the mouse is stationary (black trace) and while the mouse is locomoting (yellow trace and blue trace for the Positive and Negative group respectively). Both the Positive and the Negative group showed an excitatory response to the visual stimulus (Positive group: T1: 0.013 ± 0.006 ΔF/F_0_, T2: 0.266 ± 0.038 ΔF/F_0_, T3: 0.194 ± 0.045 ΔF/F_0_; Negative group: T1: −0.004 ± 0.009 ΔF/F_0_, T2: 0.171 ± 0.049 ΔF/F_0_, T3: 0.124 ± 0.041 ΔF/F_0_; Figure 5C,D). In the Positive group, locomotion had an excitatory effect both early in the visual stimulus presentation and late (T1: 0.112 ± 0.034 ΔF/F_0_, T2: 0.706 ± 0.172 ΔF/F_0_, T3: 0.539 ± 0.095 ΔF/F_0_; Fig 5D). In the Negative group, however, locomotion did not seem to have an excitatory effect on pre-synaptic neural activity during visual stimulation (T1: −0.080 ± 0.031 ΔF/F_0_, T2: 0.099 ± 0.087 ΔF/F_0_, T3: 0.175 ± 0.100 ΔF/F_0_; Figure 5D).

**Figure 5.**
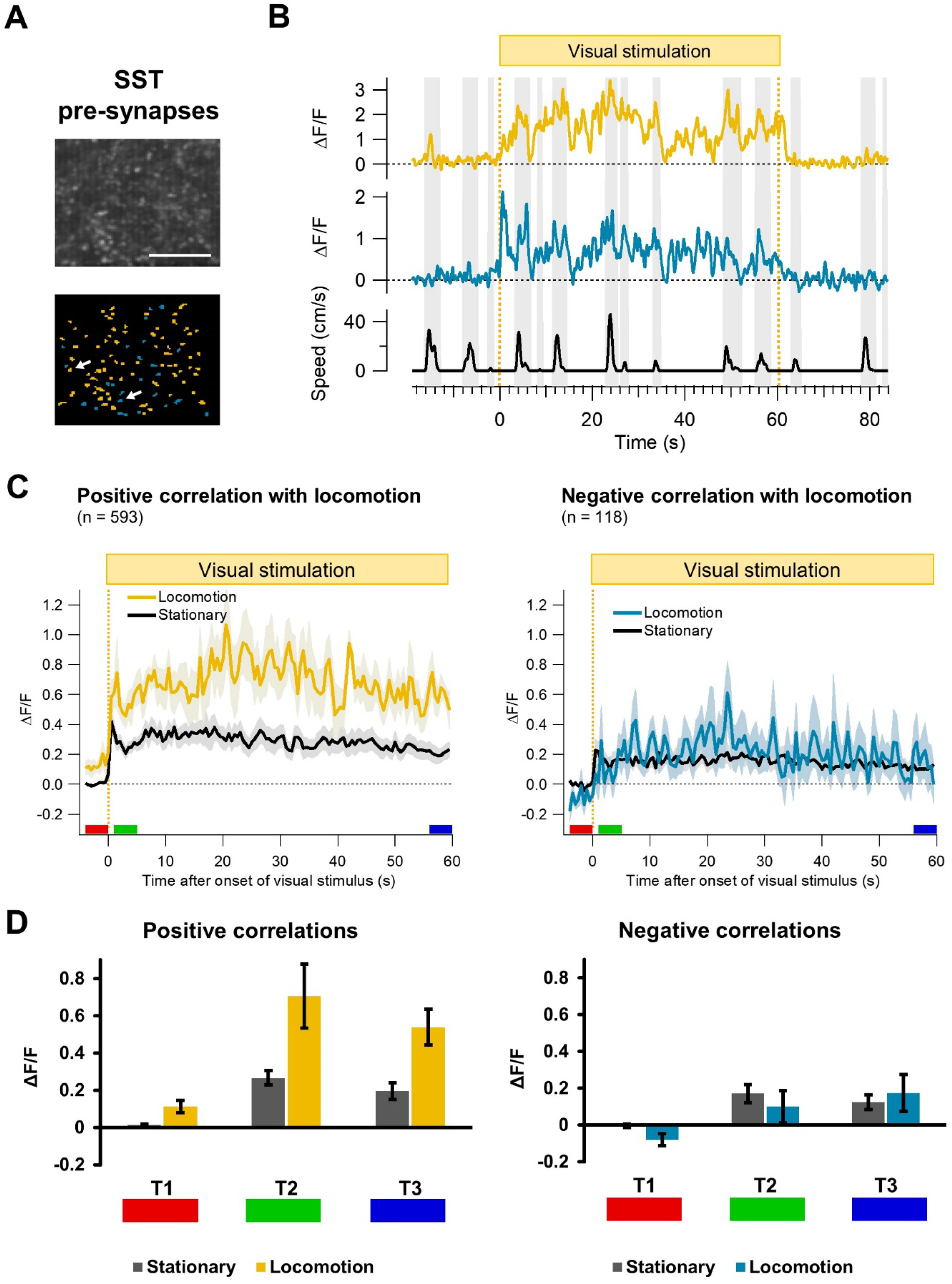
The effect of prolonged visual stimulation, with and without locomotion, on the pre-synaptic activity of SST-positive interneurons in V1 layer 2/3. **(A)** An example of a field of view (top) and the ROIs corresponding to the synapses (bottom). The ROIs showing a positive correlation with locomotion are in yellow and the ROIs showing negative correlations are in blue. Scale bar represents 20 μm. **(B)** An example of a fluorescence timeseries extracted from a ROI with a positive correlation with locomotion (yellow timeseries) and a ROI with a negative correlation (blue timeseries). The visual stimulation is indicated at the top, with the yellow dotted lines indicating the start and end of the presentation. The two ROIs are indicated in **A** by white arrows. The black trace represents the locomotion speed in cm/s. **(C)** The average somatic activity before and during prolonged visual stimulation. Synapses were separated into a group containing synapses that showed positive correlations with locomotion with p-values < 0.05 after FDR correction (Positive group) and a group representing negative correlations with p-values < 0.05 after FDR correction (Negative group). The x-axis represents the time relative to the onset of the visual stimulus and the y-axis represents the fluorescence in ΔF/F_0_. The black trace represents the synaptic activity while the mouse is stationary and the yellow (left; Positive group) and blue traces (right; Negative group) during bouts of locomotion. The light area around the traces represents SEM. **(D)** The average activity was measured at three timepoints (indicated in **C**): before onset of visual stimulation (T1, red, −4-0 sec.), early during visual stimulation (T2, green, 1-5 sec.) and after almost a minute of visual stimulation (T3, blue, 56-60 sec.). The dark grey bars represent neural activity during stationary periods and the yellow (left) and blue (right) bars activity during periods of locomotion. Error bars represent SEM.

#### Responses were generally larger in the Positive group than the Negative group

To test for significant differences between the groups and conditions, we performed a 2×3×2 mixed ANOVA. There was a significant main effect of Locomotion F(1,10)=17.533, p=0.002, n^2^- _p_=0.637, Power=0.964, Time F(2,20)=12.614, p=0.000268, n^2^_p_=0.558, Power=0.990 and Correlation group F(1,10)=15.010, p=0.003, n^2^_p_=0.600, Power=0.936.

There was no interaction between Time and Correlation group F(2,20)=2.680, p=0.093, n^2^- _p_=0.211, Power=0.470 and no three-way interaction F(2,20)=0.970, p=0.396, n^2^_p_=0.088, Power=0.195. However, the main effects were confounded by significant interactions between Locomotion and Correlation group F(1,10)=15.200, p=0.003, n^2^_p_=0.603, Power=0.939 and Locomotion and Time F(2,10)=4.235, p=0.029, n^2^_p_=0.298, Power=0.673. To determine whether each Correlation group differed from each other, independent samples T-tests were conducted separately for each Locomotion condition. Equality of variance was evaluated using Levene’s Test, which found that the assumption was not violated for either condition. The Correlation groups did not differ in the Stationary condition t(23)=1.794, p=0.086, but mean fluorescence levels were found to be significantly higher in the Positive group than the Negative group t(21)=4.065, p=0.001.

#### Visual responses in the two Correlation groups were affected by locomotion in different ways

Because the main effects were confounded, we could not draw any conclusions regarding the hypotheses. We therefore performed a series of Wilcoxon matched-pairs Signed-Rank Tests. First, we addressed the hypothesis that both Correlation groups show excitatory responses to visual stimulation both at timepoints 2 and 3 and for both Locomotion conditions. Timepoints 2 and 3 were tested against timepoint 1 for all conditions (8 tests). As we performed 12 tests in total (including 4 for the next hypothesis, see next paragraph), the p-values were adjusted to correct for multiple comparisons using the Benjamini and Hochberg False Discovery Rate method (Benjamini and Hochberg, 1995). In the Stationary condition, both timepoints 2 and 3 are significantly different from timepoint 1 for both Correlation groups (Positive group: T2 vs T1: mean difference=0.253, p=0.000244, q=0.00293; T3 vs T1: mean difference=0.181, p=0.000732, q=0.00439; Negative group: T2 vs T1: mean difference=0.175, p=0.00488, q=0.0117; T3 vs T1: mean difference=0.128, p=0.00929, q=0.01856). During the Locomotion condition, the Positive group showed a significant response to the visual stimulus for both Timepoints 2 and 3 (Positive group: T2 vs T1: mean difference=0.591, p=0.000977, q=0.00293; T3 vs T1: mean difference=0.427, p=0.0156, q=0.0267). However, no significant differences were found for the Negative group (Negative group: T2 vs T1: mean difference=0.163, p=0.313, q=0.341; T3 vs T1: mean difference=0.265, p=0.0781, q=0.104). This confirms the hypothesis that locomotion significantly increases visual responses at timepoints 2 and 3 for the Positive group and the Stationary condition of the Negative group. However, the hypothesis needs to be rejected for the Locomotion condition of the Negative group.

#### Locomotion increases the response to visual stimulation in the Positive group, but not in the Negative group

To address the second hypothesis that locomotion increases the level of fluorescence during visual stimulation in both Correlation groups, we performed 4 additional Wilcoxon matched-pairs Signed-Rank Tests. This time we tested the Locomotion condition against the Stationary condition for both Timepoints 2 and 3 and for both the Positive and Negative group. For the Positive group, a significant increase was found for both T2 and T3 (T2: Locomotion vs Stationary: mean difference=0.412, p=0.000977, q=0.00391; T3: Locomotion vs Stationary: mean difference=0.269, p=0.0156, q=0.0234), confirming the hypothesis for this group. No significant effect of locomotion during visual stimulation was found for the Negative group, however, resulting in the rejection of the hypothesis for this group (T2: Locomotion vs Stationary: mean difference=-0.00323, p=0.945, q=0.945; T2: Locomotion vs Stationary: mean difference=0.0.0650, p=0.313, q=0.341).

### The origin of the Negative group of VIP-synapses

Both for the SST- and VIP-neurons in Layer 2/3 of V1 a difference was observed in the correlations with locomotion between the soma and the pre-synapses. In SST-somas the Positive group comprised 56% and the Negative group 44% of total significant correlations. For the SST-synapses, however, the Positive group accounted for 83% of significantly correlated synapses, while the Negative group accounted for only 17%. A possible explanation for this could be that positively correlated somas give rise to the majority of pre-synaptic boutons.

#### The small number of VIP-somas showing negative correlations with locomotion do not account for the Negative group of VIP-synapses

The VIP-neurons show a more pronounced dichotomy between the somatic and pre-synaptic neural activity. Among VIP-somas, only 6% of significant correlations with locomotion were negative with the remaining 94% showing positive correlations. For VIP-synapses on the other hand, 60% were negatively correlated and 40% positively correlated with locomotion. We considered the possibility that the small group of negatively correlated somas gives rise to the Negative group of pre-synaptic boutons. VIP-synapses from the Positive group, as well as the VIP-somas, showed a pronounced increase in neural activity in response to locomotion after prolonged visual stimulation, while this feature was absent in the Negative group of VIP-synapses (Figure 3D). We therefore tested if this feature was also absent in the VIP-somas that showed negative correlations with locomotion. However, we found that the average response to locomotion late in the visual stimulation for these somas was 0.280 ± 0.033 ΔF/F_0_. This value was similar to the average response for the total population of somas (0.244 ± 0.034 ΔF/F_0_; Figure 1E) and suggests that the negatively correlated somas do not represent a population related to the Negative group of VIP-synapses.

#### V1 Layer 4 VIP-somas showed primarily positive correlations with locomotion

The concentration of VIP-interneurons is highest in layer 2/3 (Ji et al., 2015; Pakan et al., 2016). Furthermore, Karnani et al. (2016) have shown that the axons of VIP-interneurons are relatively localised to the soma. Taken together this suggests that most VIP-synapses in layer 2/3 are originating from layer 2/3 somas. However, we cannot exclude that the Negative group of VIP- synapses originates from somas in deeper layers. To address this, we collected somatic data from VIP-neurons from V1 layer 4 and calculated their correlation with locomotion. We were able to get clear responses from somas at these depths, showing that we did not reach the limitation of the imaging capabilities from deeper tissues. 70 out of 130 somas were significantly positively correlated with locomotion, while only 3 showed a significant negative correlation (N=3; Figure 6A).

**Figure 6.**
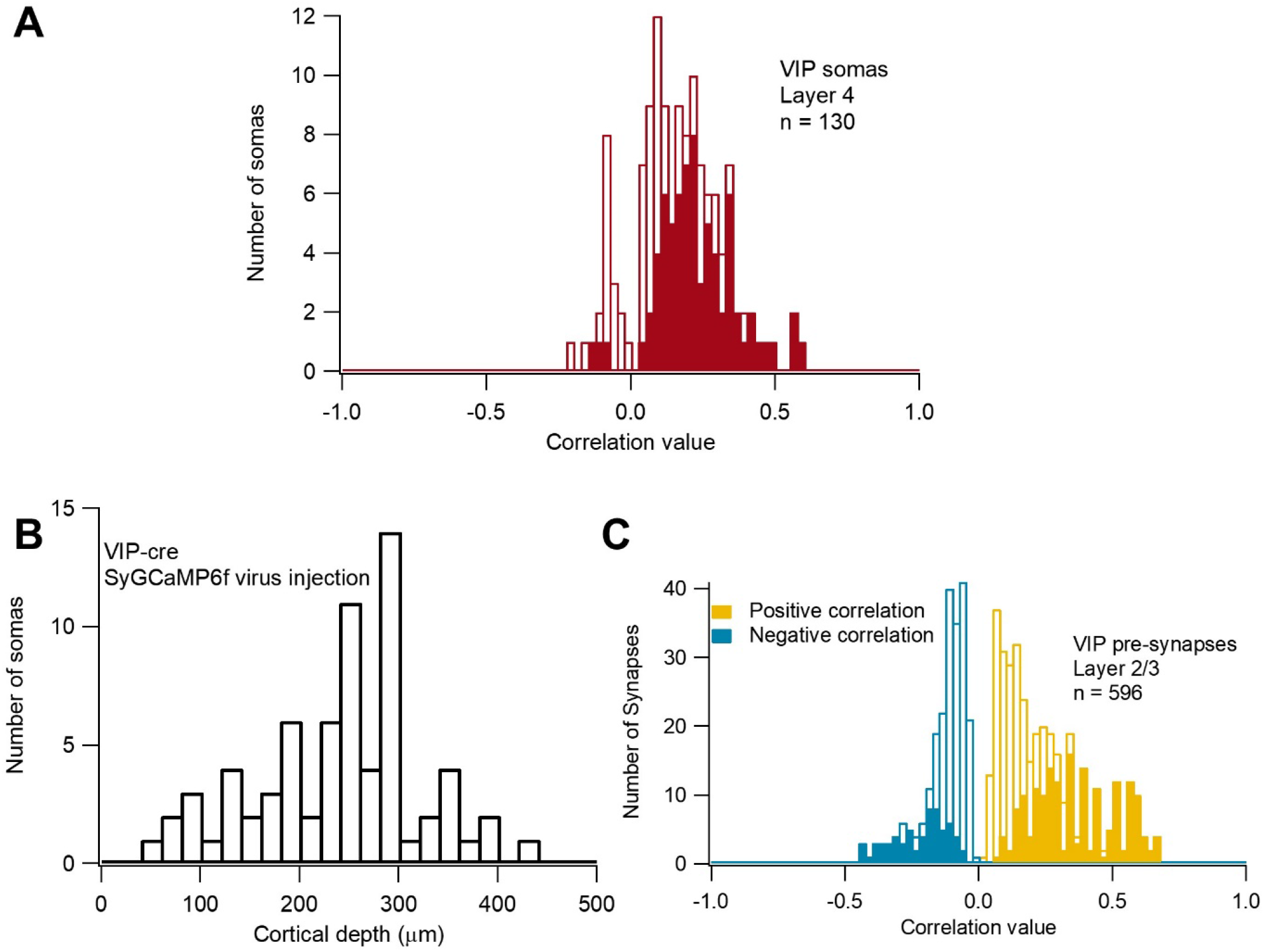
When only recording from VIP-synapses originating from V1 layer 2/3 somas, synapses that are positively correlated with locomotion outnumber synapses that show negative correlations. **(A)** Histogram of VIP-positive somas and their correlation with locomotion in V1 layer 4. n=130, recorded from 3 mice. **(B)** Flexed SyGCaMP6f virus was injected in V1 layer 2/3 of VIP-cre mice. A histogram was constructed plotting the number of somas (y-axis) and the cortical depth (μm) at which they were located. The majority of somas were located in layer 2/3 (a depth of 180-300 μm). **(C)** Histogram of VIP-synapses and their correlation value with locomotion. The filled bars represent correlations with p-values < 0.05 after FDR correction. The total number of synapses was 596, recorded from 3 mice.

#### Injection of flexed SyGCaMP6f virus in VIP-cre mice revealed a sub-group of VIP-synapses that originate solely from layer 2/3 somas

Next, we performed a viral injection of a flexed SyGCaMP6f virus in a VIP-cre driver line, localised to V1 layer 2/3. To make sure that the injection was indeed confined to layer 2/3 we made use of an artefact that resulted from the strong expression of the virus. Whereas expression of the protein is mostly confined to the pre-synaptic boutons as expected, a lower level of expression is visible in the soma. By making a z-stack of layer 1 down to layer 4 we could construct a histogram showing the number of somas present at different depths (Figure 6B). These results confirm that the injection was limited to layer 2/3 with only few cells visible in layer 4, but not deeper.

Once we were confident that the pre-synapses we recorded from were originating from V1 layer 2/3, we used the visual stimulation protocol used earlier (60 sec. visual stimulation/30 sec. grey screen; 5-10 repetitions; full field drifting Gabor grating, 315°, contrast=1, spatial frequency=0.04 cpd, temporal frequency=2Hz) to collect a dataset with the same 2×3×2 mixed design as before (see methods).

First, we calculated the Spearman correlation values for all VIP-synapses and constructed a histogram (Figure 6C). 176 out of 596 pre-synapses showed a significant positive correlation with locomotion, while only 64 synapses showed a negative correlation (N=3; Figure 6C). This was comparable to the balance found by Ryan et al. (2020) who had also made use of viral injections. To summarise, in the cross-bred mice the Negative group is larger than the Positive group (Negative:60%, Positive:40%), while in the virus-injected mice the Positive group is larger ((Negative:27%, Positive:73%). Taken together, these results suggest that only a subset of the Negative synapses in V1 layer 2/3 originate from somas within this area.

We then focussed on the effect of locomotion during prolonged visual stimulation (Figure 7A,B) and hypothesised that the results would be similar to the VIP:SyGCaMP5 cross-bred mice for both Correlation groups. To do this we conducted a 2×3×2 mixed ANOVA.

**Figure 7.**
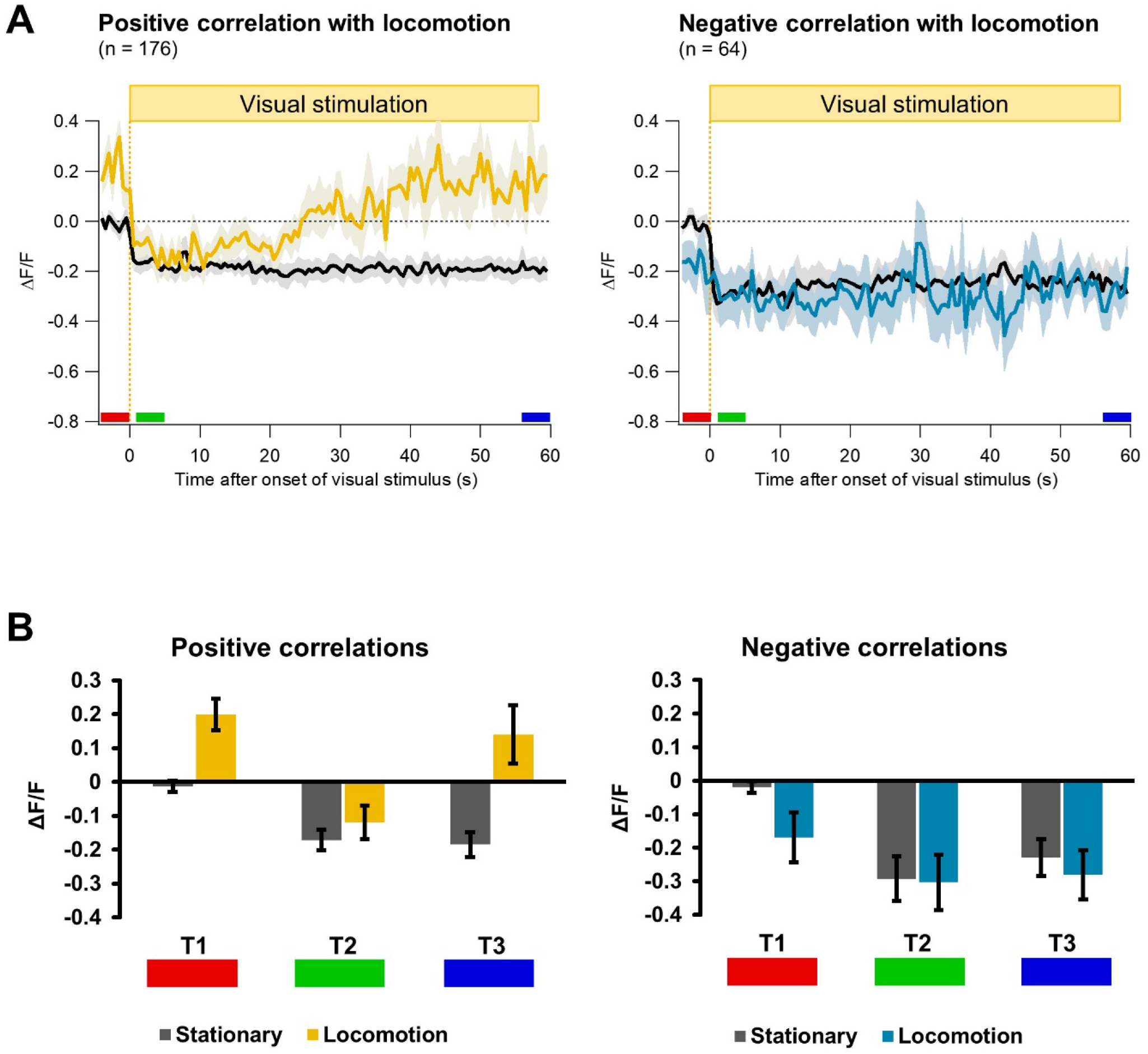
**(A)** The average synaptic activity before and during prolonged visual stimulation. Synapses were separated into a group containing synapses that showed significant positive correlations with locomotion (Positive group) and a group representing significant negative correlations (Negative group). The x-axis represents the time relative to the onset of the visual stimulus and the y-axis represents the fluorescence in ΔF/F_0_. The black trace represents the synaptic activity while the mouse is stationary and the yellow (left; Positive group) and blue traces (right; Negative group) while the mouse is in locomotion. The light area around the traces represents SEM. **(B)** The average activity was measured at three timepoints (indicated in **A**): before onset of visual stimulation (T1, red, −4-0 sec.), early during visual stimulation (T1, green, 1-5 sec.) and after almost a minute of visual stimulation (T3, blue, 56-60 sec.). The dark grey bars represent neural activity during stationary periods and the yellow (left) and blue (right) bars activity during periods of locomotion. Error bars represent SEM.

There was a significant main effect of Locomotion F(1,9)=5.391, p=0.045, n^2^_p_=0.375, Power=0.545, Time F(2,18)=16.939, p=0.000073, n^2^_p_=0.653, Power=0.999 and Correlation group F(1,9)=9.027, p=0.015, n^2^_p_=0.501, Power=0.762.

There was no interaction between Time and Correlation group F(2,18)=0.931, p=0.419, n^2^- _p_=0.092, Power=0.183 or Locomotion and Time F(2,9)=1.752, p=0202, n^2^_p_=0.163, Power=0.318, but there was a significant interaction between Locomotion and Correlation group F(1,9)=38.054, p=0.000165, n^2^_p_=0.809, Power=1. However, the main effects and 2-way interactions were confounded by a three-way interaction between Locomotion, Time and Correlation group F(2,18)=4.290, p=0.030, n^2^_p_=0.323, Power=0.671.

To further elucidate the 3-way interaction, 6 between group t-tests were conducted to determine where significant differences lay between the Positive and Negative group for each timepoint and Locomotion condition separately. To correct for multiple comparisons the alpha level was adjusted using the Bonferroni correction which resulted in an alpha of 0.05/6=0.008. There were no significant differences found between the Correlation groups in the Stationary condition for timepoint 1 t(15)=0.246, p=0.80, timepoint 2 t(13)=1.737, p=0.106, and timepoint 3 t(15)=0.685, p=0.504. For the Locomotion condition, no significant difference between the Correlation groups was found at timepoint 2 t(13)=1.931, p=0.068. However, significant differences were found at timepoint 1 t(11)=4.339, p=0.001 and timepoint 3 t(15)=3.675, p=0.002.

As we found a three-way interaction that was absent in the VIP:SyGCaMP5 crossbred mice the hypothesis was rejected. This finding further suggests that the population of VIP-synapses recorded in the virus-injected mice is different from the population recorded in the cross-bred mice.

## Discussion

In this study we have shown that the true output of VIP- and SST-interneurons is not fairly reflected by somatic activity. By comparing the neural activity at the level of the soma and the pre-synapses, we have identified a few crucial differences. The vast majority of VIP-somas are positively correlated with locomotion, whereas in the VIP-synapses we see a split between positive and negative correlations. SST-somas show a split between positive and negative correlations that is balanced slightly in favour of the positive correlations. However, in SST-synapses the positive correlations far outnumber the negative correlations. Furthermore, the response of VIP-neurons to visual stimulation is much more pronounced in VIP-synapses than VIP-somas.

VIP-interneurons are part of a disinhibitory pathway, increasing the activity of pyramidal neurons by supressing SST-positive neurons that in turn inhibit excitatory neurons as well as VIP-interneurons (Pfeffer et al., 2013; Pi et al., 2013; Fu et al., 2014; Pakan et al., 2016; Karnani et al., 2016; Dipoppa et al., 2018; Millman et al., 2020). VIP-and SST-interneurons receive cholinergic input and in this way respond to changes in behavioural states like locomotion (Picciotto et al., 2012; Alitto and Dan, 2013; Fu et al., 2014; Chen et al., 2015; Pakan et al., 2016). Here we show that, in the absence of visual stimulation, a subset of VIP-synapses increases their activation in response to locomotion. However, this increase in activity is much reduced or absent early during visual stimulation. A type of adaptation seems to then take place as the excitatory effect of locomotion slowly increases over the course of about a minute. This suggests that VIP-neurons integrate visual responses and behavioural states like locomotion in a time-dependent manner. A possible reason for this could be that a new visual stimulation is initially given priority as it may be behaviourally important, and the effect of locomotion is therefore reduced. An interesting finding was that when the mice were not exposed to the visual stimulation for at least 24 hours, the first response to the presentation was consistently excitatory (Supplementary Figure 2) instead of suppressive. It may be important that this response to such a (relatively) novel or unexpected stimulation is not obscured by locomotion. When the visual stimulus persists and becomes less relevant, the effect of locomotion becomes increasingly excitatory again. The VIP-synapses that are negatively correlated with locomotion do not show a large effect of locomotion during visual stimulation. They may fulfil a type of gain control by balancing excitation/inhibition or, more accurately, levels of inhibition (Xue et al., 2014; Millman et al., 2020).

Interestingly, the slow increase in locomotion-induced activity during prolonged full field visual stimulation in the Positive group of VIP-synapses is not reflected in the SST-synapses. This is surprising as SST-interneurons are thought to be part of the aforementioned disinhibitory pathway, receiving inhibitory inputs only from VIP-interneurons. As suggested before by Pakan et al. (2016) and Dipoppa et al. (2018), the VIP-SST-Pyramidal cell disinhibitory pathway may be more complex than we previously thought.

We compared layer 2/3 VIP-output activity in the VIP:SyGCaMP5 crossbred mouse with the VIP-cre mouse that has received an injection of flexed SyGCaMP6f virus localised to V1 layer 2/3. There was a clear difference in the ratio between synapses showing a positive and negative correlation with locomotion. Although still substantial, the portion of negatively correlated synapses in the VIP-cre mice with viral injections was a factor of 1.8 smaller than in the

VIP:SyGCaMP5 cross-bred mice. This suggests that part of the VIP-synapses within V1 layer 2/3 originated from somewhere else. Whereas the population recorded from in the cross-bred mice contained all pre-synapses present within the field of view, the population recorded from in the virus-injected mice only included pre-synapses originating in V1 layer 2/3. This was further supported by the mixed ANOVA analysis showing a three-way interaction only in the boutons from the virus-injected mice, suggesting that different populations were represented in these studies.

This difference in populations will affect the decision of whether to use cross-bred mice or virus-injected mice in a particular study. If interested in the general VIP-output in a certain area and layer, the advice would be to use the VIP:SyGCaMP cross-bred mice. However, if your research question focusses more specifically on the VIP-output originating from a certain area and layer, using flexed SyGCaMP6f virus injections in VIP-cre driver mice would be the better choice.

It seems feasible that the imbalance in ratio positive and negative correlations with locomotion for the SST-pre-synapses as compared to the SST-somas could be caused by differences in axonal arborization in SST-neuron sub-types. In this scenario, the positive group of somas may consist of an SST sub-type that gives rise to more axonal boutons than the negative group. However, while this is potentially viable for the SST-neurons, this explanation is less applicable for VIP-neurons for 2 reasons. Firstly, the group of negative somas was very small, suggesting they may have been statistical false positives. Secondly, the effect of locomotion late in the visual response of the negative soma group was very similar to that of the positive group. Another possible explanation for the dichotomy between the somatic responses and pre-synaptic responses in VIP-neurons may be a type of axonal modulation. It would be valuable for future studies to look for the presence of pre-synaptic or axonal inhibitory receptors on VIP-neurons. Cholinergic terminals from the forebrain could, for example, act on inhibitory muscarinic receptors (Felder, 1995) or corelease of GABA could trigger axonal GABA receptors (Trigo et al., 2008).

In conclusion, our results suggest that neural activity in the somas of VIP- and SST-interneurons, measured using calcium imaging, does not reflect the output of the cell accurately. Given these findings, it is therefore important to take this into account when constructing diagrams of cortical microcircuits and to wherever possible make use of synaptic recordings alongside somatic recordings.

## Author Contributions

RV has designed the study, collected the data on VIP-interneurons and written the first draft of the manuscript. LSM and RV have performed the data analysis and written the manuscript.

## Acknowledgements

We would like to thank Dr Anurag Pandey for collecting data for this study from somatostatin-expressing interneurons and for his helpful comments on the manuscript. We would also like to thank Drs Maarten Kamermans, Marcus Howlett and Christian Wozny for their helpful discussions and advice on the methods and analysis.

