## Supplementary material for "The output of interneurons in the primary visual cortex is best reflected by pre-synaptic activity, not somatic activity"

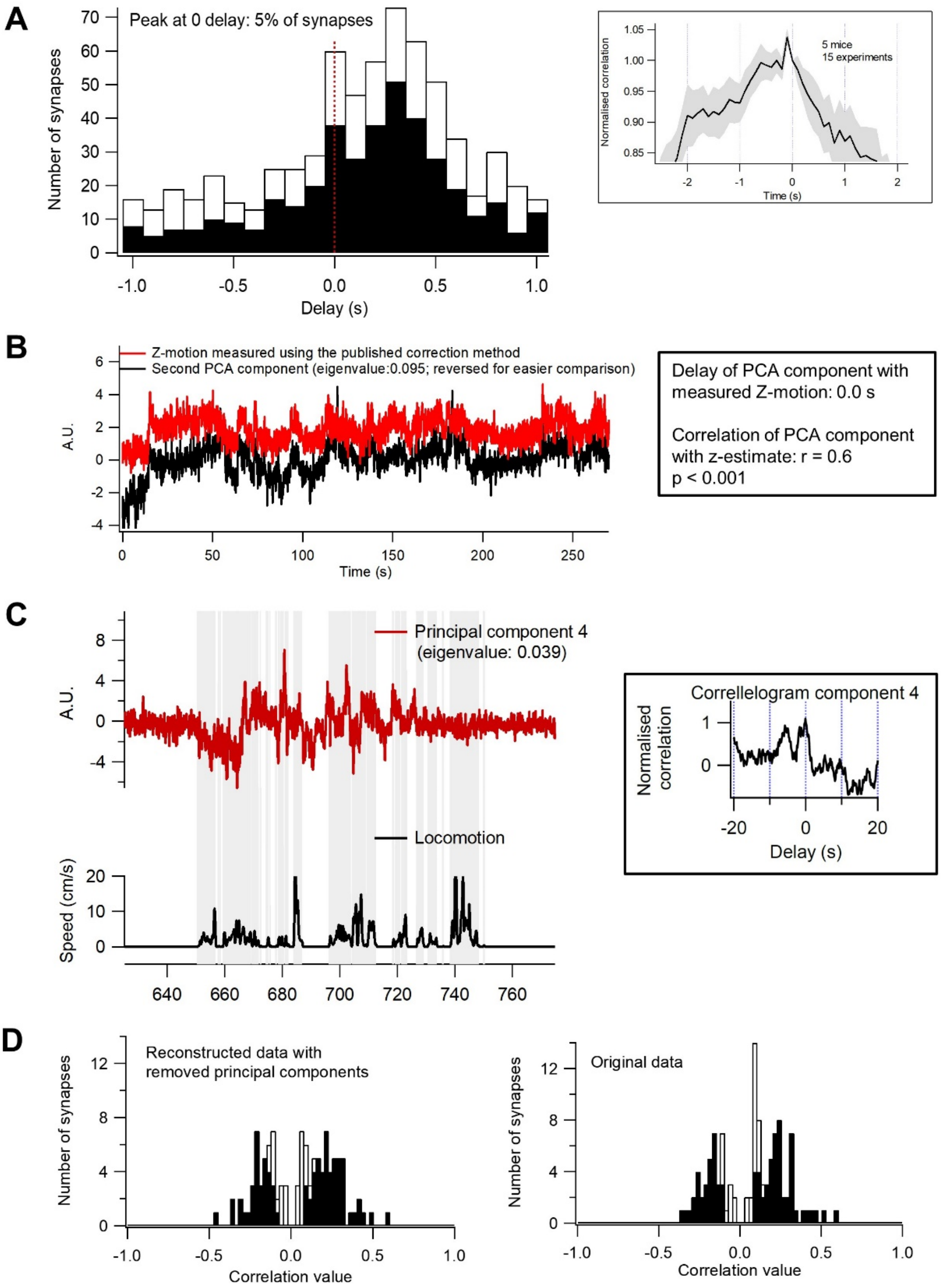

**Supplementary Figure 1**

**Removing the z-drift artefact using Principal Component Analysis**

A) The delay for all VIP pre-synapses in relation to locomotion. Locomotion-induced neural activity is delayed due to the multi-synaptic pathways involved. A histogram is shown with the delay in seconds on the x axis and the number of synapses on the y-axis. The filled bars represent the synapses showing a positive correlation with locomotion. The peak is at a delay of 300 ms. However, there is a second peak: 5% of the synapses show a delay of 0 ± 50 ms, suggesting that there is a small artefact present in the data. We set out to test whether this may be the z-motion artefact. The data has been corrected for x-y drift by using the ImageRegistration operation in Igor pro, which returns a timeseries representing the drift. Any x-y drift will potentially be accompanied by z-drift. The drift time-series was then used to measure the delay with locomotion by cross-correlating the two time-series (right graph), which showed a peak at −50 ms. This suggests that the z-drift has a delay of −50 ms.

B) A method that is commonly used in fields dealing with signal detection (e.g. fMRI and ERG) to address artefacts is Principal Component Analysis (PCA). This technique uses orthogonal transformation to convert the data into principal components that are linearly uncorrelated. Since the neural activity shows a delay with locomotion of around 300 ms and the z-motion artefact shows a delay of around 0, this technique can be used to identify the artefact. To test this, we used a dataset from the study by Ryan et al., (2020) for which the z-motion artefact was measured by making use of acquiring a reference volume before data collection. Blood vessels filled with dextran conjugated fluorescent dye were used as an anatomical marker and recorded in a second channel in addition to recording calcium signals (see Ryan et al., 2020 for a detailed explanation). The z-motion is plotted as the red trace. We then applied PCA to the calcium signal timeseries using Python 2.7 and calculated the delay with locomotion of all components using Igor pro. Components were identified as representing the artefact if their delay with locomotion was 0 ± 100 ms. The black trace represents the component identified as artefact with the highest eigenvalue, which for this dataset from Ryan et al. (2020) was the second component (we inverted it to make it easier to compare with the measured z-motion artefact). A significant correlation was found between the z-motion and the second principal component: r(2698)=0.60 (p= 5.521e-259), N=2699.

C) An example of the artefact component from one of the VIP-synapse recordings in this paper (only a section from the full timeseries is shown). The artefact in this experiment was found to be the 4$^{th}$ principal component with an eigenvalue of 0.039. The red trace is principal component 4 and the black trace is the locomotion speed readout. On the right a correlogram of the 4$^{th}$ component with locomotion shows the delay of zero.

D) After identifying the artefact components, the data was reconstructed with these components removed. On the left a histogram is shown of the locomotion correlation values calculated from the reconstructed data. On the right the histogram from the original data is shown.

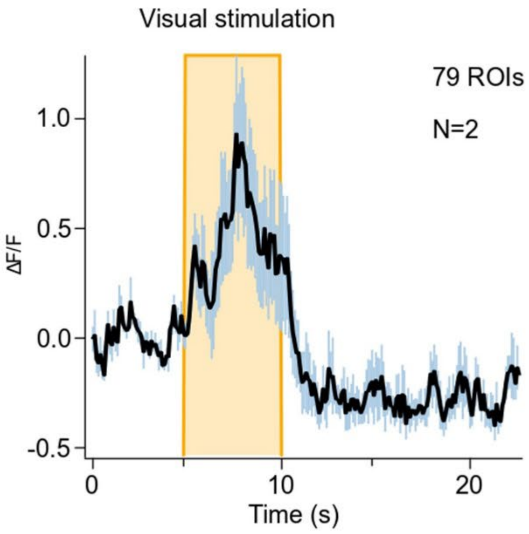


**Supplementary Figure 2**

If mice were not exposed to the visual stimulation (full field drifting Gabor grating, 315°, contrast=1, 0.04 cpd, 2Hz) over a period of 24 hours or more, the first response was consistently excitatory. Average timeseries from 79 VIP-synapses, recorded from 2 mice. Grey area represents SEM.

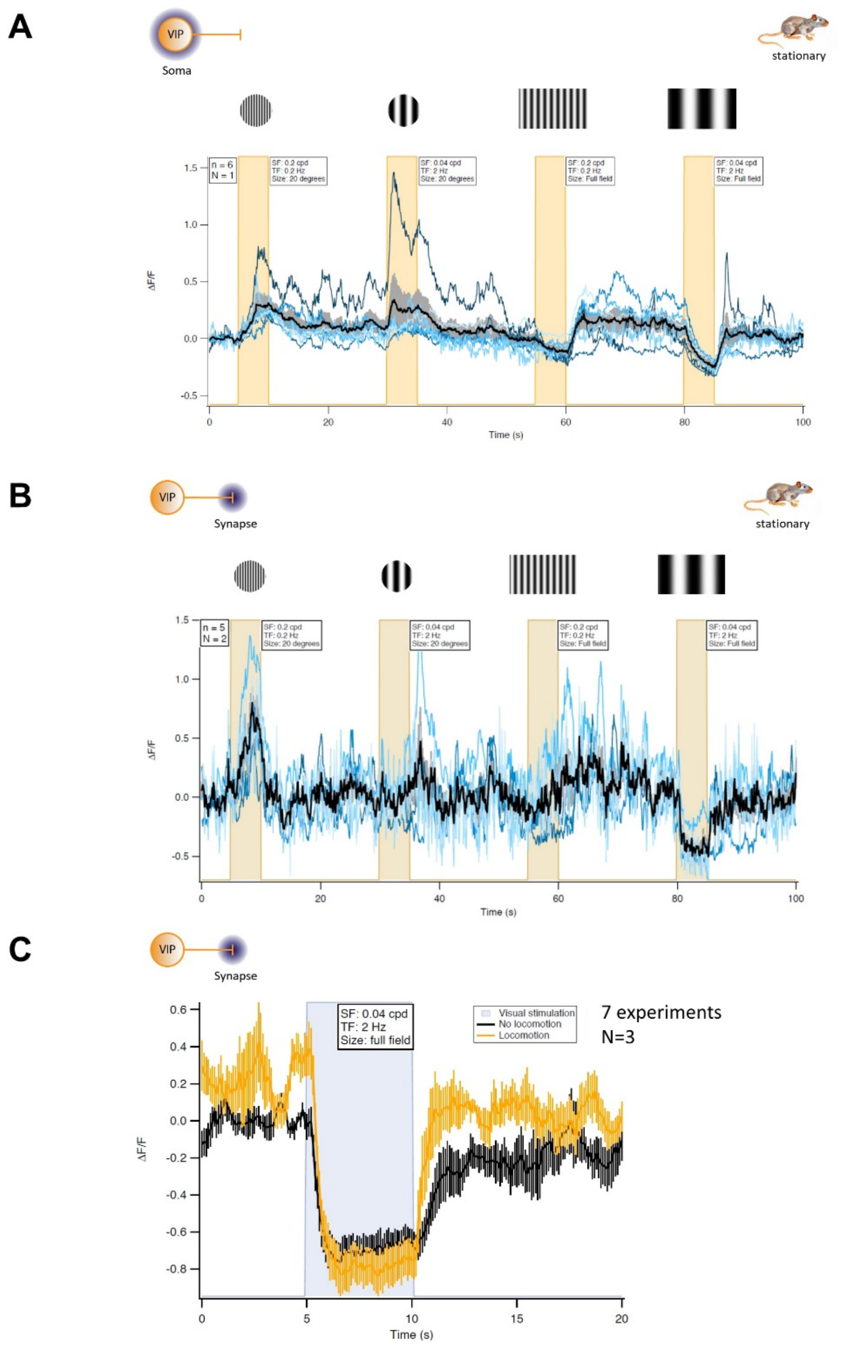


**Supplementary Figure 3**

Preliminary data showing responses in VIP-somas and VIP-synapses to 20 degree and full field stimuli. **(A)** Somatic activity in response to two 20-degree visual stimuli and two full field stimuli with 2 sets of condition: 0.2 cpd, 0.2 Hz and 0.04 cpd, 2 Hz. The set of 4 visual stimulations was repeated 10 times. The black trace represents the average of 6 somas from 1 mouse, the grey area the SEM and the blue traces are the individual measurements. **(B)** The same protocol was used. Recorded from 5 VIP-synapses from 2 VIP:SyGCaMP mice. **(C)** The average response to a 5 second full field visual stimulation (0.04 cpd, 2Hz) from 7 experiments recorded from 3 VIP:SyGCaMP mice. The black timeseries represents stationary mice and the yellow timeseries mice in locomotion.
